# A targeted plasma-proteomic axis separates autoimmune thyroid disease from growth-hormone deficiency

**DOI:** 10.64898/2026.07.27.741048

**Authors:** Eunjeong Han, Jaeho Ji, Yunho Choi, Jumin Park, Heejin Lee, Sujin Park, Ahrum Son, Sukdong Yoo, Chong Kun Cheon, Hyunsoo Kim

**Affiliations:** Graduate School of Life Sciences, Chungnam National University, Daejeon 34134, Republic of Korea; Department of Convergent Bioscience and Informatics, Chungnam National University, Daejeon 34134, Republic of Korea; Institute of Molecular Life Sciences, Chungnam National University, Daejeon 34134, Republic of Korea; Graduate School of Medical Science, University of Ulsan, Ulsan 44610, Republic of Korea; Department of Pediatrics, Pusan National University School of Medicine, Pusan National University Children’s Hospital, Yangsan 50612, Republic of Korea; Research Institute for Convergence of Biomedical Science and Technology, Pusan National University Yangsan Hospital, Yangsan 50612, Republic of Korea; Department of Bio-AI Convergence, Chungnam National University, Daejeon 34134, Republic of Korea; Protein Design Institute, Chungnam National University, Daejeon 34134, Republic of Korea; SCICS, Daejeon 34134, Republic of Korea

**Keywords:** Targeted proteomics, Plasma biomarkers, Autoimmune thyroid disease, Growth-hormone deficiency, Machine-learning classification

## Abstract

Targeted mass spectrometry (multiple-reaction monitoring, MRM) enables reproducible, multiplexed quantification of plasma proteins, but whether a fixed targeted panel can resolve endocrine disorders with overlapping systemic features is unknown. We analyzed a 256-protein targeted panel (1,894 peptides; 3,790 transitions) quantified in 57 participants spanning autoimmune thyroid disease (Hashimoto’s thyroiditis, n=7; Graves’ disease, n=5) and growth-hormone deficiency (GHD; partial, n=26; complete, n=19). Protein abundances were obtained by transition summation, log2 transformation, and per-sample median normalization. We applied unsupervised analysis (PCA, PERMANOVA), differential expression (limma), an ordered severity-trend test, and leave-one-out cross-validated classification with feature selection performed strictly inside each fold. All 256 proteins were quantified in every sample (median inter-sample r=0.875). PC1 (36% variance) separated autoimmune thyroid disease from GHD (p=0.016), whereas the global four-group structure was not significant (PERMANOVA p=0.17). No protein reached FDR<0.05, but the autoimmune-versus-GHD contrast was strongly enriched for low p-values (26 proteins at p<0.05; binomial p=5.4×10^−4^). The signal was biologically coherent: immunoglobulin/B-cell-receptor proteins, including CD79A, were lower, whereas proteasome subunits (PSMC5, PSMC3) and the NF-κB subunit RELA were higher in autoimmune disease. A cross-validated classifier separated the two classes (AUC 0.72; permutation p=0.05; eight proteins selected in all folds), whereas GHD severity was not predictable (AUC 0.31). A fixed 256-protein targeted panel reproducibly captures an immunoglobulin/B-cell-receptor and proteasome/NF-κB axis that distinguishes autoimmune thyroid disease from GHD but cannot resolve within-class severity.

## Introduction

Endocrine disorders are among the most common chronic conditions worldwide, yet many remain difficult to diagnose and to distinguish from one another because they share nonspecific systemic features and rely on assays with well-recognized analytical and interpretive limitations [1, 2]. Two disorders illustrate this problem from opposite ends of the endocrine spectrum: autoimmune thyroid disease and growth-hormone deficiency (GHD). Both are prevalent, both present with overlapping constitutional symptoms — fatigue, altered metabolism, and impaired growth or development — and both are diagnosed today through combinations of hormone measurements, provocative testing, and autoantibody assays that are imperfect, operator- or assay-dependent, and in some cases invasive [2–4]. A minimally invasive, reproducible, blood-based molecular readout able to support the differential diagnosis of endocrine disease would therefore address a genuine and still-unmet clinical need.

Autoimmune thyroid disease — comprising Hashimoto’s thyroiditis and Graves’ disease — is the most common organ-specific autoimmune disorder, affecting roughly 5% or more of the population in iodine-replete regions and predominantly women [3, 5]. It arises from a breakdown of self-tolerance to thyroid antigens, with autoreactive T- and B-lymphocyte activation, thyroid-directed autoantibody production, and lymphocytic infiltration of the gland [6, 7]. B cells and antibody-secreting plasma cells are central effectors; Graves’ disease in particular is regarded as an archetypal B-cell-mediated disorder in which autoreactive B cells expand and secrete pathogenic anti-thyrotropin-receptor antibodies [1, 8]. These adaptive-immune programs are transcriptionally coordinated by the nuclear factor-κB (NF-κB) family, whose activity is increased in thyroid autoimmunity and whose canonical RelA-containing dimers drive inflammatory gene expression and B-cell survival [9, 10]. Clinically, however, the autoantibodies that define the disease are an incomplete readout: their titers correlate only loosely with disease activity, a substantial minority of patients are seronegative, and the assays do not capture the broader systemic immune state that accompanies thyroid autoimmunity [11, 12]. Markers that reflect this systemic footprint, rather than a single antigen-specific antibody, could add diagnostic and mechanistic value.

Growth-hormone deficiency, by contrast, is an endocrine-metabolic disorder of the somatotropic axis; unlike the antibody-driven autoimmune process above, its diagnosis rests on auxology together with low insulin-like growth factor-1 (IGF-1) and subnormal responses to growth-hormone stimulation testing, and it is graded clinically along a partial-to-complete severity continuum [2, 13]. Yet the diagnostic pathway for GHD is among the least satisfying in endocrinology. Growth-hormone stimulation tests depend on arbitrary and shifting cut-offs, are poorly reproducible within and between individuals, are confounded by assay heterogeneity, adiposity, and pubertal status, and typically require two provocative challenges to reach a still-uncertain conclusion [2, 4, 14]. IGF-1, the most widely used single biomarker, has limited sensitivity and specificity, particularly near the boundary separating partial from complete deficiency [13, 14]. A reproducible circulating signature that tracked the somatotropic axis would therefore ease a diagnostic process that is currently invasive, costly, and imprecise. Together, these two conditions — an antibody-driven autoimmune disorder at one pole and a metabolic hormone-axis deficiency at the other — make the pairing an informative test of what a fixed targeted panel can, and cannot, capture.

The plasma proteome is an attractive substrate for such markers because blood is the most accessible clinical specimen and continuously integrates proteins secreted, shed, or leaked from virtually every tissue and cell type in the body [15–17]. In practice, however, the panel of proteins used routinely in the clinic remains small, and new plasma biomarkers reach validation slowly, chiefly because untargeted discovery pipelines are difficult to reproduce across laboratories and instruments, so promising candidates frequently fail to transfer to independent cohorts [18, 19]. Targeted mass spectrometry — multiple-reaction monitoring (MRM) — was developed to close this reproducibility gap by quantifying a predefined set of peptides through defined precursor–fragment transitions, yielding assays that transfer across sites and instrument platforms with high precision and quantitative accuracy [20–23]. A fixed, validated panel of dozens to hundreds of proteins can then be interrogated with supervised machine learning to assign a sample to a diagnostic category. This paradigm has been most successful for conditions that leave a strong, coordinated systemic footprint on the circulating proteome — for example, plasma-proteomic classifiers that separate clinically overlapping fibrotic lung diseases otherwise difficult to distinguish [24]. Whether the same fixed-panel strategy can resolve endocrine disorders, whose circulating perturbation may be smaller and more diffuse, is far less clear, and defining where such panels succeed and, equally important, where they fail is a prerequisite for any clinical deployment.

Here we analyze a uniformly processed targeted plasma-proteomics dataset spanning four diagnoses within these two classes and ask three questions. First, does unsupervised proteome structure track diagnostic class? Second, which proteins, if any, discriminate the classes, and what biology do they encode? Third, can a rigorously cross-validated classifier assign disease class and within-class severity? To avoid the optimistic bias that arises when feature selection is performed on the full dataset before validation, all feature ranking, scaling, and model fitting were confined to the training partition of each cross-validation fold.

We find that a fixed 256-protein panel reproducibly captures a plasma-proteomic axis — anchored by an immunoglobulin/B-cell-receptor module and a proteasome/NF-κB module — that separates autoimmune thyroid disease from GHD, while it cannot resolve within-class GHD severity. By reporting both the reproducible signal and its explicit limits, this study calibrates realistic expectations for targeted plasma-proteomic classification in endocrinology and nominates concrete, testable markers for clinical validation.

## Materials and Methods

### Study population and cohort design

This study included a total of 57 pediatric patients presenting with endocrine disorders. The study population was stratified into four distinct clinical diagnostic groups based on their clinical manifestations, laboratory findings, and diagnostic evaluations: Group 1 (Hashimoto’s Thyroiditis, HT, n = 7), Group 2 (Graves’ Disease, GD, n = 5), Group 3 (Partial Growth Hormone Deficiency, pGHD, n = 26), and Group 4 (Complete Growth Hormone Deficiency, cGHD, n = 19). Baseline characteristics including gestational age (GA) at birth, birth weight (Bwt), age at diagnosis, and anthropometric measurements (current height, current weight, and body mass index [BMI]) were systematically collected at the time of evaluation. Clinical signs, such as goiter, exophthalmos, and family history of thyroid or endocrine diseases, were evaluated. Diagnostic laboratory assessments for thyroid function and autoantibody profiles (TSH, Free T4, TSH receptor antibody [TR-Ab], Anti-TPO Ab, and Anti-TG Ab) were measured for Groups 1 and 2. For growth hormone deficiency cohorts (Groups 3 and 4), serum insulin-like growth factor-1 (IGF-1), insulin-like growth factor-binding protein 3 (IGF-BP3), basal growth hormone (GH), and peak GH levels following stimulation testing were analyzed. Diagnostic assignments were established clinically and provided with the dataset (**Table 1; Table S1**). Two orthogonal analytical axes were defined a priori: a between-class comparison (autoimmune thyroid disease versus GHD) and within-class comparisons (Hashimoto’s versus Graves’ disease, and GHD partial versus complete, the latter treated as an ordered severity axis).

**Table 1.**
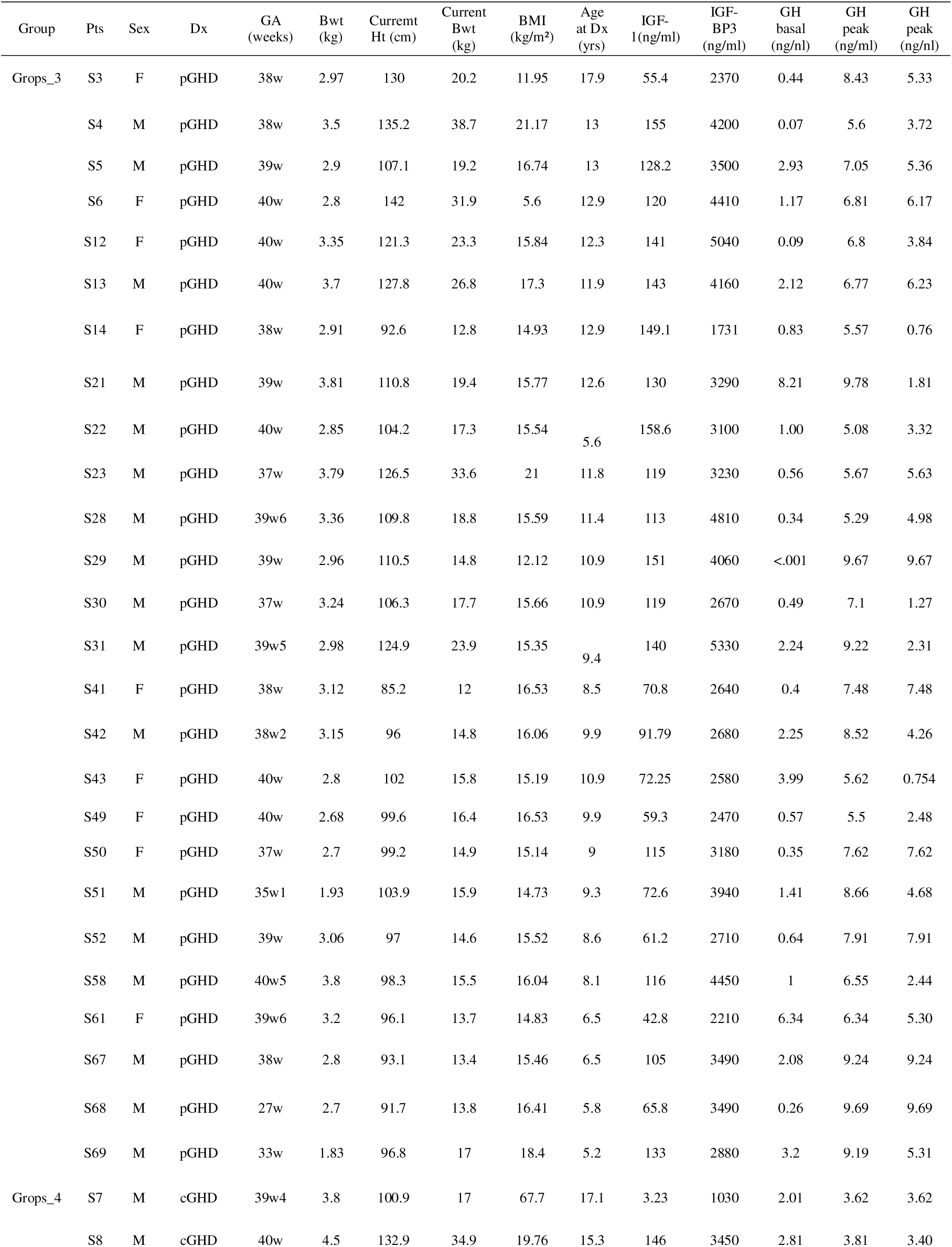
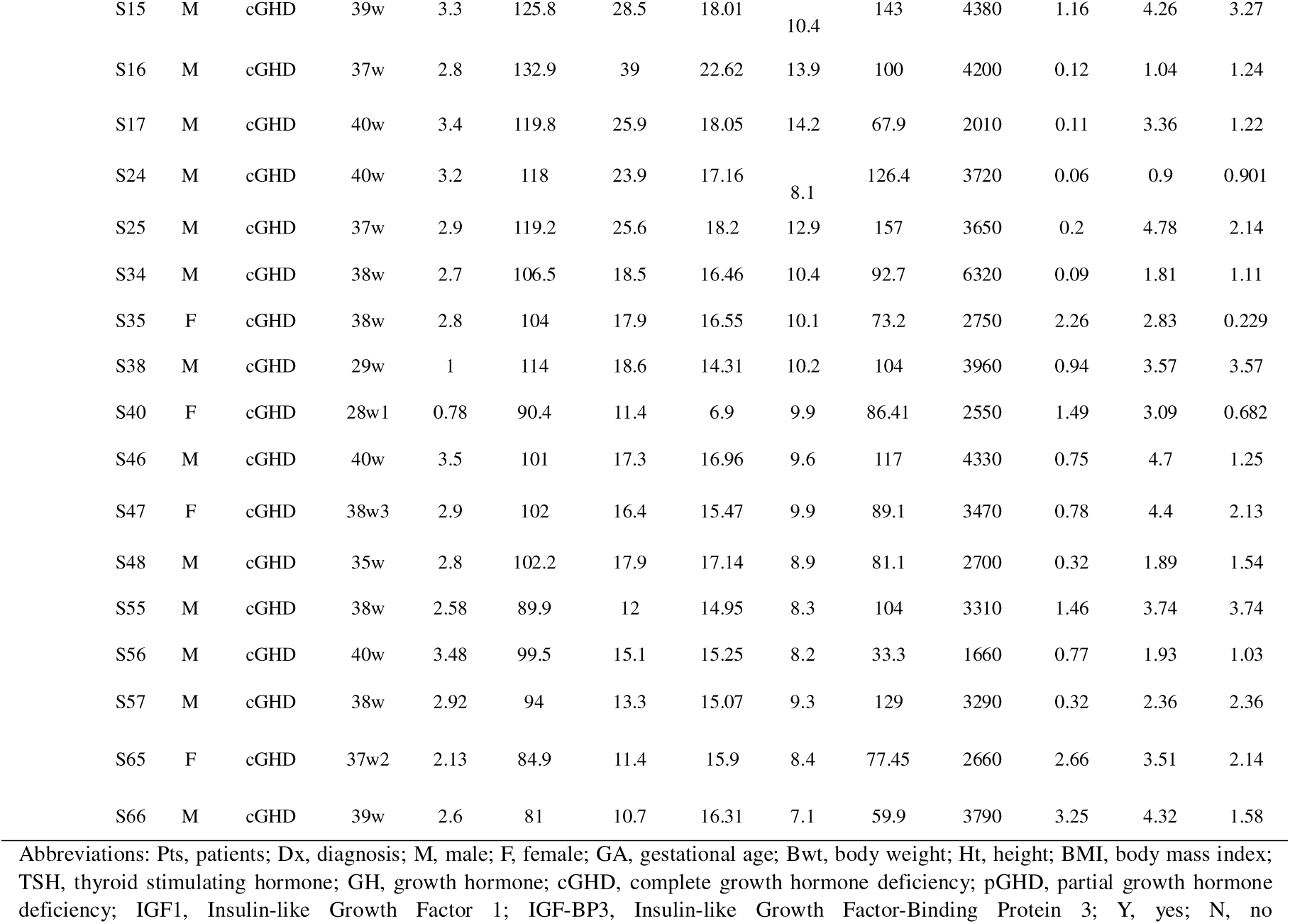
Baseline Demographic and Clinical Characteristics of the Study Population.

### Ethics approval and consent

The study was conducted in accordance with the principles of the Declaration of Helsinki and was approved by the Institutional Review Board of Pusan National University Yangsan Hospital (IRB approval no. [11-2024-046]). Written informed consent was obtained from all adult participants; for participants who were minors, written informed consent was obtained from a parent or legal guardian together with age-appropriate assent. All plasma samples were de-identified before proteomic analysis.

### Protein digestion

Plasma samples were prepared for LC-MS/MS-based quantitative proteomic analysis by enzymatic digestion. Each tube, containing 100 μg of protein, was first treated to reduce disulfide bonds using 2% sodium deoxycholate (SDC) and 10 mM tris-(2-carboxyethyl)-phosphine hydrochloride (TCEP) at 60 for 60 minutes. To block free cysteines, 20 mM iodoacetamide (IAA) was added and the samples were incubated in the dark at 25 for 30 minutes. The protein solution was then diluted to a final concentration of 20 mM ammonium bicarbonate (ABC) with distilled water. Trypsin (Promega, Sequencing Grade Modified, 20 μg) was added at a 1:50 enzyme-to-protein ratio and digestion proceeded overnight at 37□. The reaction was quenched with 1% formic acid. Finally, 5 μL of a 6 x 5 LC-MS/MS Peptide Reference mix (Promega) was added to the digested samples, which were stored at -20 until ready for analysis.

### Mass spectrometry

Digested plasma was analyzed on an Agilent 1290 Infinity II UHPLC coupled to an Agilent 6495D triple-quadrupole mass spectrometer with electrospray ionization. Mobile phases were (A) 0.1% formic acid in water and (B) 0.1% formic acid in acetonitrile. Detectability was first assessed on pooled plasma in unscheduled MRM mode to confirm transitions. Method building and data processing were performed in Skyline (v22.2). Decoy transitions (reversed sequences at 50% of the target-transition rate) were included to support mProphet peak scoring. Each unscheduled method contained up to 186 transitions (cycle time 1000 ms). Peaks were integrated with an mProphet scoring model retaining the top three transitions per precursor. Quantification of individual samples (n = 57) then used scheduled MRM (4-min retention window, up to 165 concurrent transitions, three injection methods; injection volume 10 µL, cycle time 1000 ms), processed in Skyline with the same mProphet model at q < 0.05. Integrated peak areas were exported for statistical analysis.

### Targeted proteomic dataset

Quantitative measurements were obtained from a targeted MRM assay and supplied in format (replicate × target × normalized peak area). Each target encoded a proteotypic peptide together with its parent protein (UniProt accession), precursor m/z, precursor charge state, and fragment-ion index. After parsing and quality filtering, the assay resolved 3,790 transitions mapping to 1,894 peptides and 256 proteins. The quantitative matrix was complete — every transition was measured in every sample, with no missing values — which removes the imputation step that is a well-recognized source of variability and bias in untargeted proteomics.

### Protein-level quantification, transformation, and normalization

Transition-level intensities were aggregated to the protein level by summation of all transitions mapping to a given protein within each sample. Summed abundances were log2-transformed after addition of an offset equal to one-half of the smallest non-zero value in the matrix, which stabilizes the variance of low-abundance measurements and avoids undefined logarithms. To correct for per-sample differences in total loading and instrument response, each sample was median-centered by subtracting its own median in log2 space. The resulting 256 × 57 protein-abundance matrix served as the input for all downstream analyses (**Table S2**).

### Quality control and unsupervised structure analysis

Data quality was assessed by three complementary metrics: detection completeness (the fraction of proteins quantified per sample), the distribution of protein abundances before and after normalization, and the inter-sample correlation structure (pairwise Pearson correlations), summarized as the median across all sample pairs. Global proteome structure was then examined without reference to diagnostic labels. Principal-component analysis (PCA) was computed on mean-centered, unit-variance-scaled abundances; separation along individual components was tested by one-way ANOVA across the four groups and by the Mann–Whitney U test for the two-class contrast. Overall multivariate group structure was tested by permutational multivariate analysis of variance (PERMANOVA; Euclidean distance, 9,999 permutations) [25] for both the four-group configuration and the GHD partial-versus-complete contrast. Agreement between unsupervised hierarchical clustering (Ward’s linkage on Euclidean distances) and the diagnostic labels was quantified by the adjusted Rand index (ARI).

### Differential abundance analysis

Differential protein abundance was assessed with the limma empirical-Bayes framework, which moderates per-protein variance estimates by borrowing information across proteins and is well suited to small-sample designs [26]. Moderated t-statistics were computed with the mean–variance trend enabled (trend=TRUE, robust=FALSE). Five contrasts were evaluated: autoimmune thyroid disease versus GHD; Hashimoto’s versus Graves’; GHD complete versus partial; and each of Hashimoto’s and Graves’ versus all remaining samples. p-values were adjusted for multiple testing across proteins within each contrast by the Benjamini–Hochberg procedure [27]. Because no protein reached a 5% false-discovery rate, we additionally tested whether the number of nominally significant proteins (p<0.05) exceeded the chance expectation using an exact binomial test, and estimated the proportion of true null hypotheses (π□) to gauge the overall strength of signal in each contrast.

### Severity-trend and disease-specific analyses

For the ordered GHD severity axis (partial → complete), monotonic dose-response trends were tested for each protein with the non-parametric Jonckheere–Terpstra test for ordered alternatives, with Benjamini–Hochberg adjustment across proteins. Disease-specific signals were explored by one-versus-rest Welch’s t-tests for each of the four diagnoses, providing a complementary view of proteins selectively elevated or reduced in a single diagnostic group.

### Cross-validated classification with in-fold feature selection

Two binary classification tasks were defined: autoimmune thyroid disease versus GHD (whole cohort, n=57) and GHD complete versus partial (n=45). Model performance was estimated by leave-one-out cross-validation. To prevent information leakage and the resulting optimistic bias in performance estimates [28], the entire modeling pipeline — univariate feature ranking and selection, standardization, and model fitting — was executed inside each training partition only; the held-out sample contributed to neither feature selection nor scaling. L2-regularized logistic regression with balanced class weights and random forests were compared across selected-feature counts (k = 10, 20, 30). Discrimination was summarized by the area under the receiver-operating-characteristic curve (AUC) computed on the pooled out-of-fold predictions, together with balanced accuracy, sensitivity, precision, and F1 score. The statistical significance of the primary AUC was assessed by a label-permutation test (200 permutations). Feature stability was quantified as the fraction of folds in which each protein was selected, providing a reproducibility-weighted view of the discriminating signature.

### Functional annotation and module definition

To interpret the discriminating signal, all 256 UniProt accessions were queried through the UniProt REST interface [29] for protein name, gene symbol, keywords, and Gene Ontology biological-process terms. Proteins were grouped into functional categories, and the leading discriminating proteins were mapped to their molecular roles to define coherent functional modules.

### Statistical software and reproducibility

Analyses were performed in Python (pandas, scikit-learn [30], statsmodels, and matplotlib) and R (limma [26]). All processing scripts, intermediate matrices, and result tables are provided as reproducible artifacts to allow independent re-execution of the full pipeline.

## Results

### Clinical Manifestation and Demographics

The demographic and baseline clinical characteristics of all participants across the four study groups are summarized in Table 1. Hashimoto’s Thyroiditis (Group 1, n = 7): comprised 5 females (71.4%) and 2 males (28.6%). The mean age at diagnosis was 8.8 ± 2.8 years (range: 5.7–12.9 years). A family history of thyroid disease was observed in 5 out of 7 patients (71.4%), while goiter was clinically present in 3 patients (42.9%). None of the patients in Group 1 presented with exophthalmos. Elevated TSH levels and anti-thyroid autoantibodies (Anti-TPO Ab and Anti-TG Ab) were notably present in this group. Graves’ Disease (Group 2, n = 5) included 4 females (80.0%) and 1 male (20.0%), with a mean age at diagnosis of 11.4 ± 4.9 years. Goiter was noted in 3 patients (60.0%), and exophthalmos was detected in 2 patients (40.0%). Three patients (60.0%) reported a positive family history. Patients in this cohort demonstrated suppressed TSH levels alongside elevated TSH receptor antibody (TR-Ab) positivity. Partial Growth Hormone Deficiency (Group 3, n = 26) cohort consisted of 16 males (61.5%) and 10 females (38.5%). The age at diagnosis ranged from 5.2 to 17.9 years. Diagnostic evaluation revealed intermediate peak GH levels following stimulation testing (mean peak GH: 4.9 ± 2.6 ng/mL, 6.7± 1.6 ng/mL, respectively). Complete Growth Hormone Deficiency (Group 4, n = 19) group was composed of 15 males (78.9%) and 4 females (21.1%), with an age range at diagnosis from 7.1 to 17.1 years. Patients with cGHD exhibited significantly lower peak GH levels compared to the pGHD group during stimulation testing (mean peak GH: 1.9 ± 1.1 ng/mL, 3.3 ± 1.2 ng/mL, respectively).

### A complete, high-quality 256-protein plasma matrix across 57 participants

The targeted panel returned a fully populated quantitative matrix: all 256 proteins were detected and quantified in all 57 samples, with no missing values across any of the 3,790 transitions. This completeness is a direct consequence of the targeted acquisition strategy and removes the missing-value imputation step that complicates untargeted proteomic analyses and can distort downstream statistics. After log2 transformation and per-sample median normalization, the offset abundance distributions became closely aligned across all samples, and the median pairwise inter-sample Pearson correlation was 0.875 (**Figure 1**). Neither detection completeness nor total per-group signal differed systematically among the four diagnostic groups, indicating that the subsequent analyses were not confounded by global differences in sample quality or loading. Together, these properties make the dataset well suited to both unsupervised structure analysis and supervised classification.

**Figure 1.**
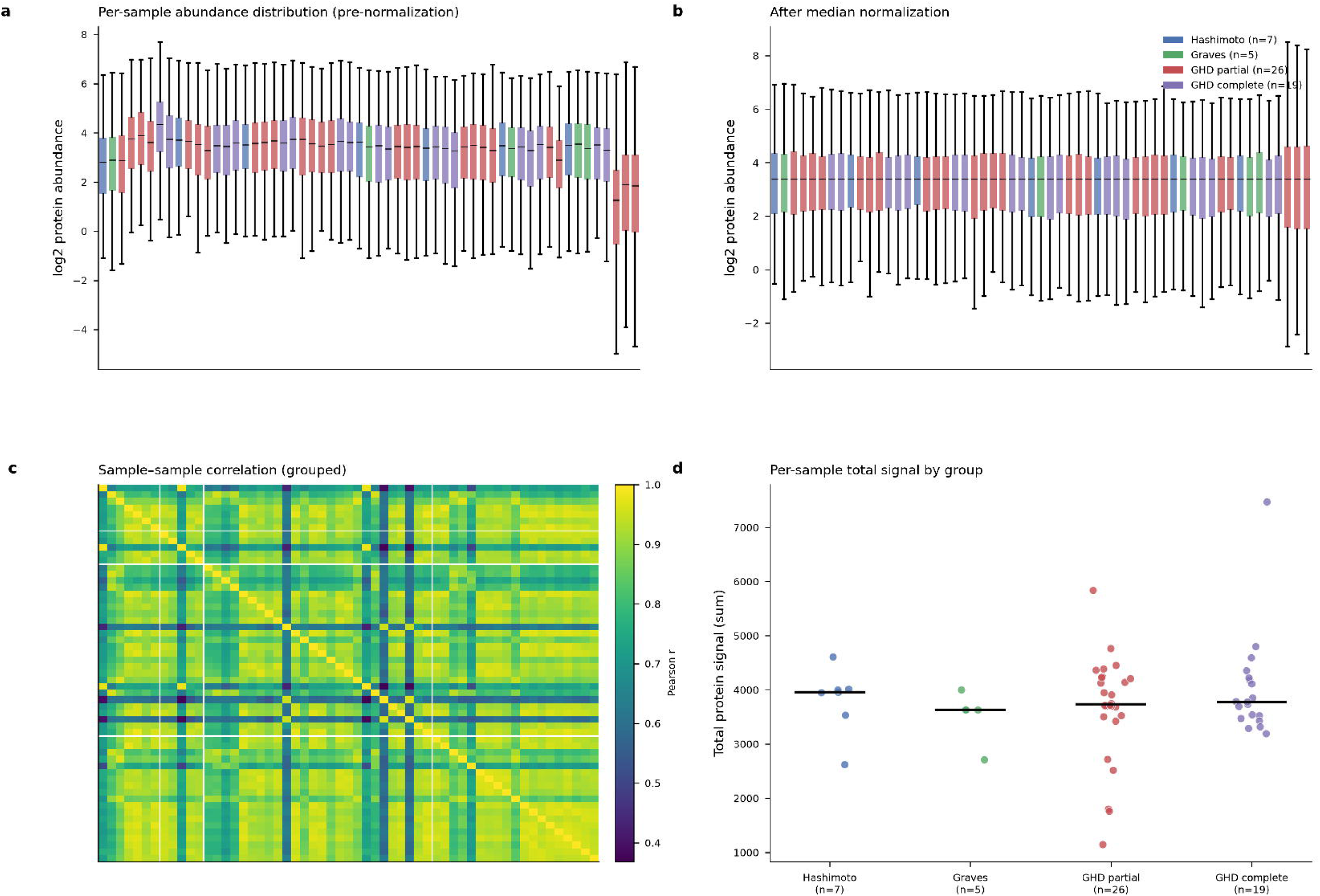
Quality control and normalization of the 256-protein targeted plasma-proteomic dataset. (a) Per-sample distributions of log2 protein abundance across the 57 samples before normalization, colored by diagnostic group (Hashimoto’s thyroiditis, n = 7; Graves’ disease, n = 5; GHD partial, n = 26; GHD complete, n = 19); box plots reveal sample-to-sample offsets in overall signal attributable to differences in loading and instrument response. (b) The same per-sample distributions after log2 transformation and per-sample median centering, which align the medians and interquartile ranges across all samples and render abundances directly comparable. (c) Heatmap of pairwise inter-sample Pearson correlation coefficients (samples ordered by group); correlations are uniformly high (median pairwise r = 0.875), with a small number of lower-correlation samples visible as darker rows and columns. (d) Per-sample total protein signal (summed abundance) by group, with group medians indicated; total signal did not differ systematically among the four groups, excluding global sample-quality or loading differences as drivers of the downstream between-class contrasts. Across all 57 samples, every one of the 256 proteins (3,790 transitions) was quantified with no missing values, so no imputation was required.

### Unsupervised proteome structure separates disease classes locally rather than globally

Principal-component analysis showed that the first principal component captured 36% of the total variance and separated the autoimmune thyroid class from GHD (Mann–Whitney p=0.016 on PC1), whereas subsequent components carried substantially less structured, group-related variance (**Figure 2**). This class-level separation was, however, not reflected in the global multivariate configuration: the four-group PERMANOVA was not significant (p=0.17), unsupervised hierarchical clustering did not recover the diagnostic labels (ARI ≈ 0), and the GHD partial-versus-complete split was likewise not separable at the whole-proteome level. The juxtaposition is itself informative. It indicates that the diagnostic information carried by this panel is local — concentrated in a limited set of proteins that load onto PC1 — rather than expressed as a diffuse, proteome-wide shift that would dominate the overall distance structure. This pattern directly motivated the protein-level differential-expression and feature-selection analyses that follow, which are designed to isolate exactly such localized signals.

**Figure 2.**
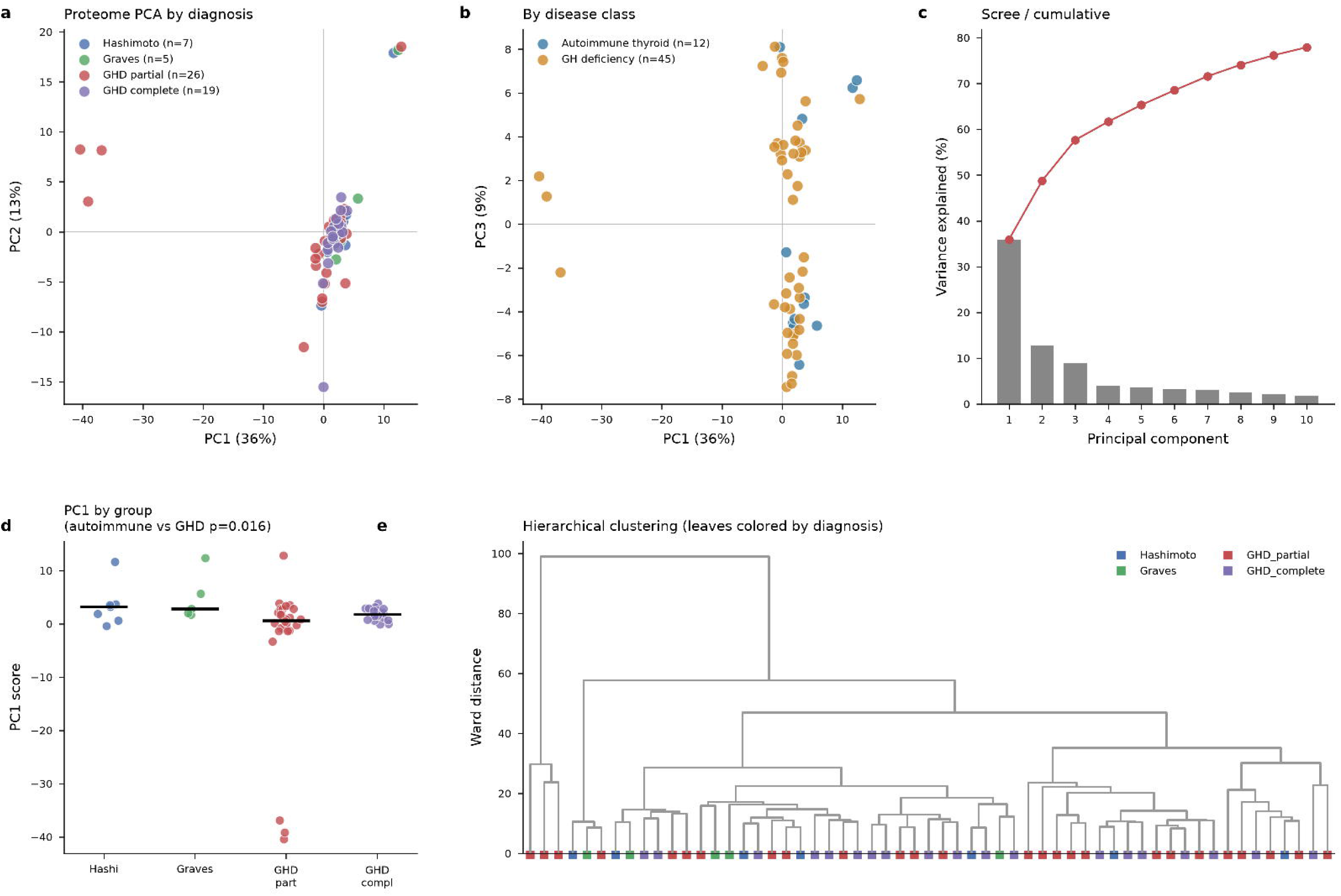
Unsupervised proteome structure across diagnostic groups. (a) Principal-component analysis (PCA) score plot of PC1 versus PC2 for the normalized 256-protein matrix, with samples colored by the four diagnoses; PC1 and PC2 explain 36% and 13% of the total variance, respectively. (b) PCA score plot of PC1 (36%) versus PC3 (9%), with samples colored by the two disease classes (autoimmune thyroid disease, n = 12; GH deficiency, n = 45). (c) Scree/cumulative plot of the variance explained by successive principal components; PC1 dominates, with a marked drop thereafter. (d) Distribution of PC1 scores by diagnostic group; PC1 separates the autoimmune thyroid class from GHD (Mann–Whitney U test, p = 0.016), whereas within-class differences along PC1 are minimal. (e) Hierarchical clustering dendrogram (Ward’s linkage on Euclidean distances) with leaves colored by diagnosis; the recovered clusters do not correspond to the diagnostic labels (adjusted Rand index ≈ 0), and, consistently, PERMANOVA (Euclidean distance, 9,999 permutations) detected no significant global four-group structure (p = 0.17). Together the panels show that diagnostic information is carried locally, by a limited set of proteins loading onto PC1, rather than as a diffuse, proteome-wide shift.

### A coordinated immunoglobulin/B-cell-receptor and proteasome/NF-*κ*B axis discriminates autoimmune thyroid disease from GHD

No single protein survived a 5% false-discovery-rate threshold in any of the five contrasts — a limitation we state plainly and attribute chiefly to the small size of the autoimmune groups. Nevertheless, the autoimmune-versus-GHD contrast carried a clear, non-random signal that was absent from the other contrasts. Twenty-six proteins reached nominal significance (p<0.05), roughly double the ∼13 expected under the global null (exact binomial p=5.4×10□□); six proteins reached FDR<0.10; and the estimated proportion of true nulls was π□ ≈ 0.62, indicating that a substantial minority of proteins genuinely differ between the classes (**Figure 3; Table S3**). By contrast, the two within-class comparisons and the disease-specific one-versus-rest tests yielded at most chance-level counts of nominally significant proteins.

**Figure 3.**
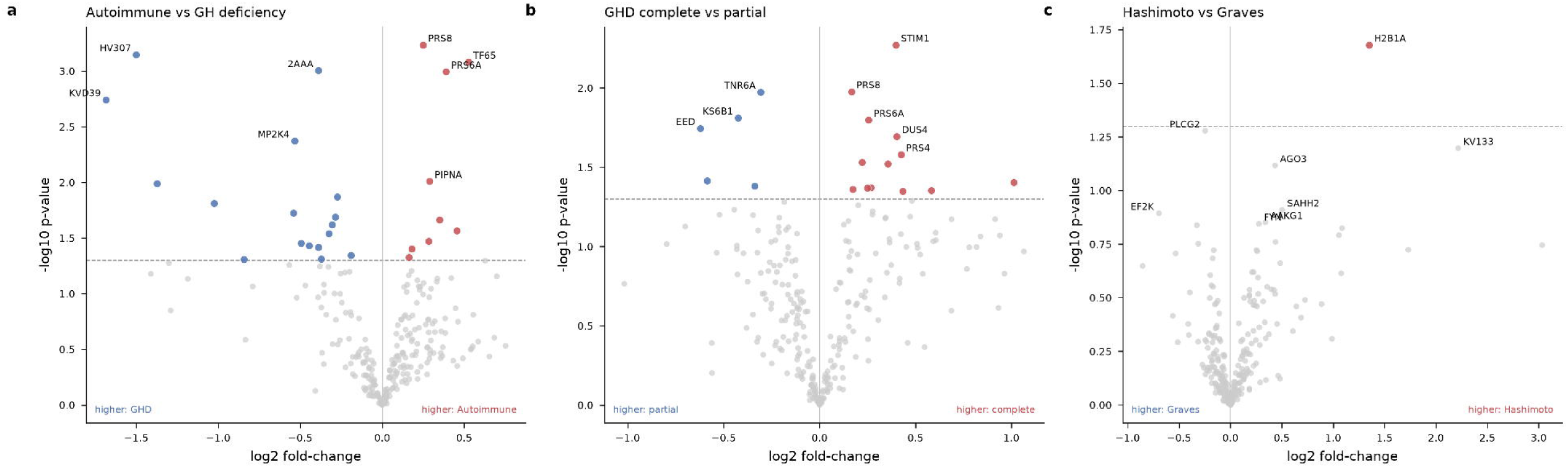
Differential protein abundance across the three diagnostic contrasts. Volcano plots of log2 fold-change (x-axis) versus −log10 nominal p-value (y-axis) from limma empirical-Bayes moderated t-tests for (a) autoimmune thyroid disease versus GH deficiency, (b) GHD complete versus partial, and (c) Hashimoto’s thyroiditis versus Graves’ disease. In each panel the dashed line marks the nominal p < 0.05 threshold, the direction of higher abundance is annotated at the lower corners, and proteins passing the threshold are colored by that direction and labeled with their assay entry name. No protein reached a 5% false-discovery rate (FDR) in any contrast. However, the autoimmune-versus-GHD contrast (a) was strongly enriched for low p-values — 26 proteins at nominal p < 0.05, roughly double the ∼13 expected by chance (exact binomial test, p = 5.4×10□□), with 6 proteins at FDR < 0.10 and an estimated true-null proportion π ≈ 0.62 — whereas neither within-class contrast (b, c) showed enrichment of low p-values beyond the chance expectation. Leading proteins in (a) include the immunoglobulin variable-region peptides HV307 (IGHV3-7) and KVD39 (IGKV1D-39), higher in GHD, and the proteasome/NF-κB proteins PRS8 (PSMC5), PRS6A (PSMC3) and TF65 (RELA), higher in the autoimmune class.

Crucially, the discriminating proteins were not a statistically arbitrary set but assembled into a coherent biological picture (**Table 2**; **Figure 4**). Immunoglobulin heavy- and kappa-variable-region peptides (IGHV3-7, IGKV1D-39, IGKV1-33) and the B-cell-receptor signaling component CD79A were consistently lower in the autoimmune thyroid class, with the largest reductions in Graves’ disease. In the opposite direction, the 26S-proteasome regulatory ATPase subunits PSMC5 and PSMC3 and the NF-κB transcription-factor subunit RELA were higher in the autoimmune class. This bidirectional, module-structured pattern — a coordinated shift in adaptive-immune/immunoglobulin proteins together with proteasome and NF-κB machinery — recapitulates the known immunobiology of autoimmune thyroid disease, in which B-cell and antibody responses against thyroid antigens are orchestrated by NF-κB-dependent transcription and sustained by proteasome-dependent antigen processing [6–10].

**Figure 4.**
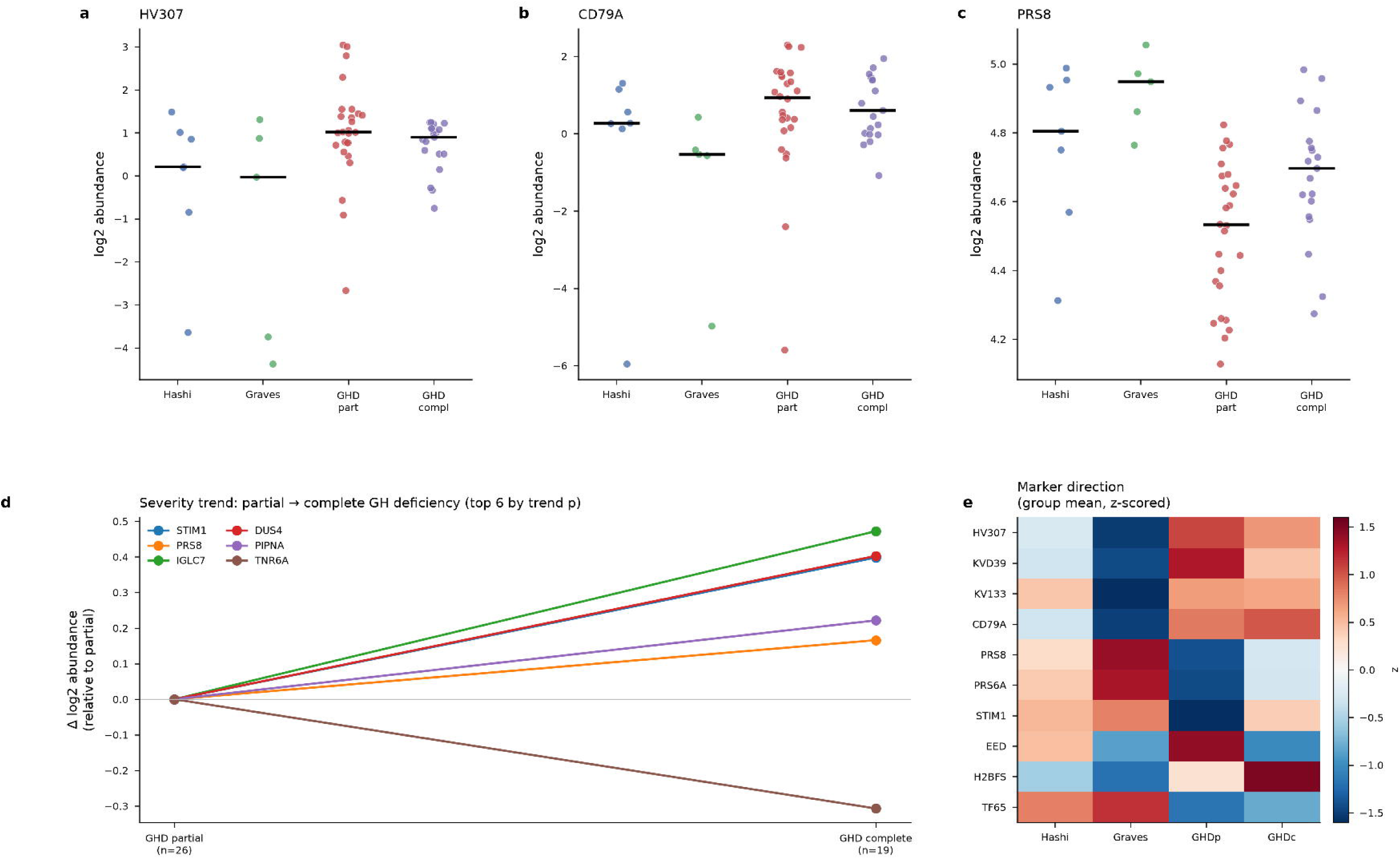
Leading discriminating proteins and growth-hormone-deficiency severity trajectories. (a–c) Per-sample log2 abundance of three representative discriminating proteins across the four diagnostic groups (horizontal bars denote group medians): (a) HV307 (IGHV3-7) and (b) CD79A, immunoglobulin/B-cell-receptor-module proteins that are lower in the autoimmune thyroid groups — most markedly in Graves’ disease — and (c) PRS8 (PSMC5), a proteasome-module protein that is higher in the autoimmune groups. (d) Severity trajectories of the six proteins with the strongest ordered partial→complete trend, plotted as the change in log2 abundance relative to GHD partial (n = 26) at GHD complete (n = 19); STIM1, PRS8 (PSMC5), IGLC7, DUS4 and PIPNA increase toward complete deficiency while TNR6A decreases, but none survived multiple-testing correction (minimum FDR ≈ 0.47). (e) Heatmap of z-scored group-mean abundance for the leading discriminating proteins across the four groups (Hashi, Graves, GHDp, GHDc), summarizing the direction and magnitude of each protein’s between-group difference. Together the panels show that the discriminating signal is biologically coherent and recapitulates the known immunobiology of autoimmune thyroid disease.

**Table 2.** Leading discriminating proteins (autoimmune thyroid disease vs growth-hormone deficiency). Group means are log2 median-normalized abundances. logFC and p from limma; severity-trend p from the Jonckheere–Terpstra test. Immunoglobulin/B-cell-receptor proteins are lower and proteasome/NF-κB proteins higher in the autoimmune class.

| Protein | Gene | UniProt | Hashimoto | Graves | GHD partial | GHD complete | log<sub>2</sub>FC (Auto vs GHD) | p (Auto vs GHD) | p (severity trend) | Direction |
| --- | --- | --- | --- | --- | --- | --- | --- | --- | --- | --- |
| HV307 | IGHV 3-7 | P01780 | -0.1 | -1.19 | 1.02 | 0.69 | -1.5 | 0.0007 | 0.1542 | Down in autoimmune |
| KVD39 | IGKV 1D-39 | P04432 | -0.53 | -1.61 | 1.02 | 0.22 | -1.68 | 0.0018 | 0.2063 | Down in autoimmune |
| KV133 | IGKV 1-33 | P01594 | -0.15 | -2.36 | 0.07 | 0.02 | -1.3 | 0.0532 | 0.9633 | Down in autoimmune |
| CD79A | CD79 A | P11912 | -0.32 | -1.21 | 0.54 | 0.67 | -1.37 | 0.0103 | 0.6624 | Down in autoimmune |
| PRS8 | PSMC 5 | P62195 | 4.76 | 4.92 | 4.51 | 4.67 | 0.25 | 0.0006 | 0.0123 | Up in autoimmune |
| PRS6A | PSMC 3 | P17980 | 4.69 | 4.9 | 4.28 | 4.53 | 0.39 | 0.001 | 0.0243 | Up in autoimmune |
| TF65 | RELA | Q04206 | 4.04 | 4.14 | 3.52 | 3.61 | 0.53 | 0.0008 | 0.2905 | Up in autoimmune |
| STIM1 | STIM 1 | Q13586 | 4.63 | 4.68 | 4.2 | 4.6 | 0.25 | 0.1016 | 0.0038 | Up in autoimmune |
| EED | EED | O75530 | 3.67 | 3.33 | 3.91 | 3.29 | -0.1 | 0.7187 | 0.0243 | Down in autoimmune |
| H2BFS | H2BC 12L | P57053 | -0.63 | -1.11 | -0.05 | 0.88 | -1.29 | 0.1415 | 0.2602 | Down in autoimmune |
| IGLC7 | IGLC 7 | A0M8Q6 | 1.13 | 1.11 | 0.5 | 0.98 | 0.38 | 0.1668 | 0.0139 | Up in autoimmune |
| TNR6A | TNRC 6A | Q8NDV7 | 5.24 | 5.27 | 5.54 | 5.23 | -0.13 | 0.316 | 0.0203 | Down in autoimmune |

### Growth-hormone-deficiency severity is not encoded in the targeted panel

We next asked whether the panel could grade GHD along its clinical severity axis. Testing each protein for a monotonic partial→complete trend identified several nominal associations (STIM1 p=0.0038, PSMC5 p=0.012, IGLC7 p=0.014), but none survived multiple-testing correction (minimum FDR ≈ 0.47; **Table S4**). The panel therefore did not provide statistically robust markers of growth-hormone-deficiency severity, a negative result that stands in deliberate contrast to the reproducible between-class signal described above.

### Cross-validated classification confirms a reproducible class axis but not severity

To integrate the localized differential signal into a single decision rule while guarding against selection bias, we trained leave-one-out classifiers with all feature selection performed strictly inside each fold. The autoimmune-versus-GHD classifier separated the two classes with an AUC of 0.72 (balanced accuracy 0.65; sensitivity 0.80; label-permutation p=0.05). Performance was stable across model families and feature-set sizes — AUC ranged from 0.67 to 0.72 for logistic regression and random forests at k = 10, 20, and 30 — and eight proteins were selected in 100% of folds, defining a compact and reproducible core signature (**Figure 5; Tables S5, S6**). In sharp contrast, the GHD complete-versus-partial classifier achieved an AUC of 0.31, below the chance expectation of 0.5 and consistent with the absence of a severity-related differential signal. The discordance between a reproducible, biologically interpretable class axis and an entirely absent severity axis is the central quantitative result of this study and defines the resolution limit of the assay in this cohort.

**Figure 5.**
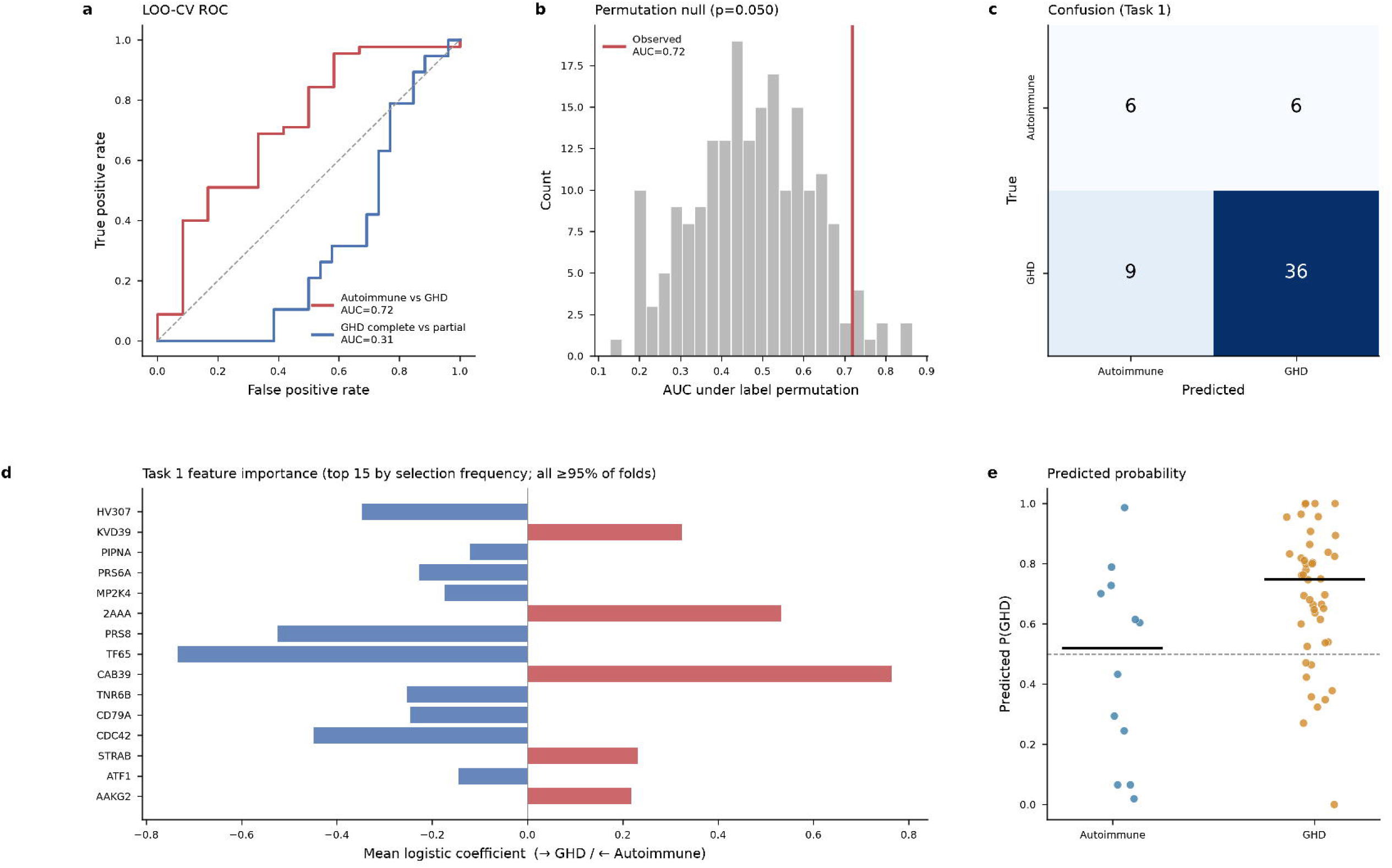
Cross-validated classification of disease class and severity. (a) Leave-one-out cross-validated receiver-operating-characteristic (ROC) curves for both classification tasks: autoimmune thyroid disease versus GHD (red; area under the curve, AUC = 0.72) and GHD complete versus partial (blue; AUC = 0.31, below the chance diagonal). For both tasks, univariate feature selection, scaling, and model fitting were performed strictly within each training fold to prevent information leakage. (b) Null distribution of AUC values for the class task under 200 label permutations; the observed AUC (0.72) lies at the upper tail (permutation p = 0.05). (c) Confusion matrix for the class task at the default operating point: of 12 autoimmune samples, 6 were correctly classified and 6 misclassified as GHD; of 45 GHD samples, 36 were correctly classified and 9 misclassified as autoimmune (balanced accuracy 0.65; GHD sensitivity 0.80). (d) The 15 proteins most frequently selected across folds (all selected in ≥ 95% of leave-one-out folds; eight in 100%), shown as their mean multivariable logistic-regression coefficient, with the sign indicating the class toward which each protein drives the prediction (positive → GHD; negative → autoimmune). The stably selected features include the immunoglobulin/B-cell-receptor and proteasome/NF-κB module members HV307, KVD39, CD79A, PRS8 (PSMC5), PRS6A (PSMC3) and TF65 (RELA). (e) Predicted probability of GHD, P(GHD), for samples of each true class (horizontal bars, group medians), showing partial but consistent separation. Class-classifier performance was stable across model families and feature-set sizes (AUC 0.67–0.72 for logistic regression and random forests at k = 10, 20, 30).

### Functional architecture of the panel constrains the detectable biology

Annotation of 254 of 256 proteins showed that the panel was weighted toward intracellular regulatory machinery: transcription regulation (86 proteins), kinase/phosphatase signaling (63), the proteasome/ubiquitin system (37), immunoglobulin/B-cell-receptor components (23), and calcium signaling (10) (**Figure 6; Table S7**). The two disease-associated modules identified above — an immunoglobulin/B-cell-receptor module and a proteasome/NF-κB module — therefore fall squarely within the regions of biology that the panel was designed to interrogate. This composition helps explain the study’s central asymmetry: a panel emphasizing immune, transcriptional, and protein-turnover machinery is well positioned to detect the adaptive-immune axis of autoimmune thyroid disease, but poorly positioned to capture the metabolic gradations that distinguish partial from complete growth-hormone deficiency. The detectable biology is thus bounded by the design of the assay itself, a consideration that should guide the composition of future targeted panels intended for endocrine classification.

**Figure 6.**
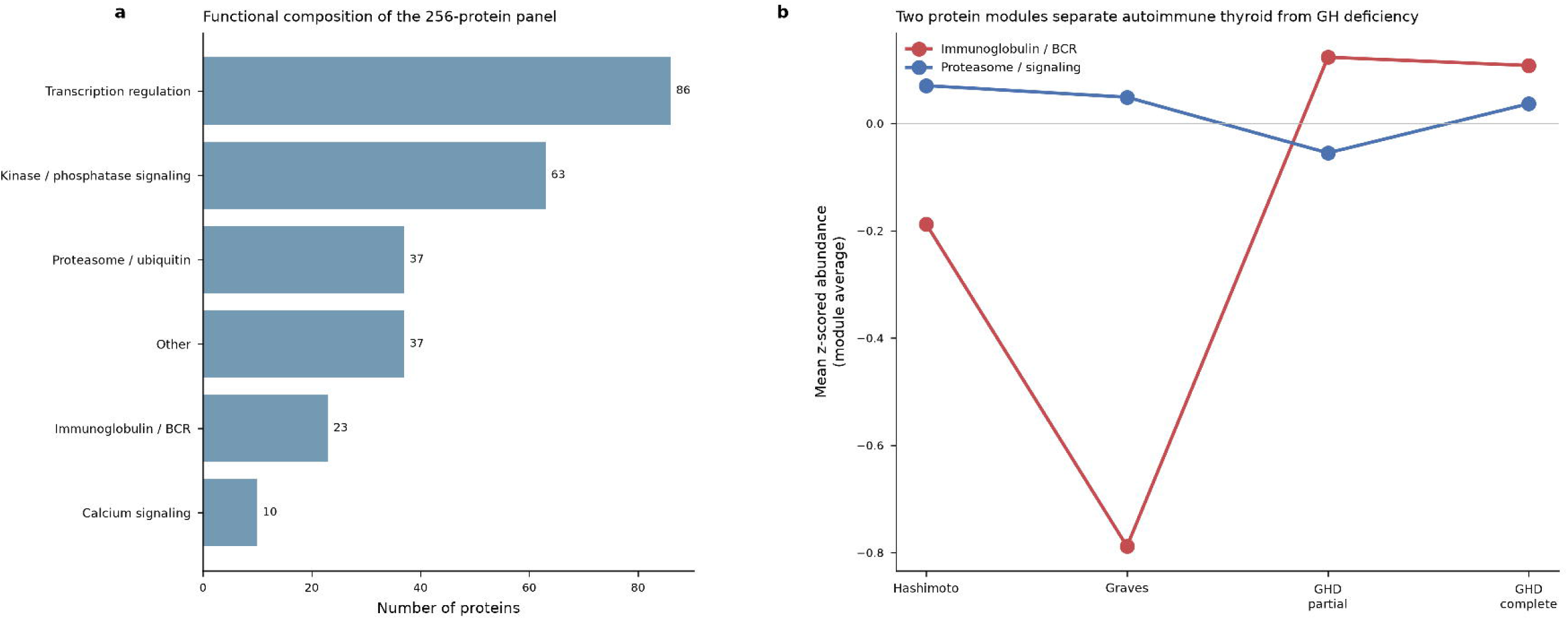
Functional architecture of the panel and the two disease-associated modules. (a) Functional composition of the 256-protein panel by UniProt keyword/Gene Ontology category: transcription regulation (86 proteins), kinase/phosphatase signaling (63), proteasome/ubiquitin (37), other (37), immunoglobulin/B-cell receptor (23), and calcium signaling (10). (b) Mean z-scored abundance (module average) of the immunoglobulin/B-cell-receptor module (red) and the proteasome/signaling module (blue) across the four diagnostic groups (Hashimoto’s, Graves’, GHD partial, GHD complete). The immunoglobulin/B-cell-receptor module is markedly reduced in the autoimmune thyroid groups — lowest in Graves’ disease — and higher in GHD, whereas the proteasome/signaling module shows the opposite, milder pattern, so that the two modules move in opposing directions between the disease classes. The panel illustrates that the discriminating modules fall within the regions of biology the panel was designed to interrogate.

## Discussion

We analyzed a 256-protein targeted plasma-proteomics dataset across four endocrine diagnoses and found a reproducible proteomic axis that separates autoimmune thyroid disease from growth-hormone deficiency. The axis is anchored by two biologically coherent modules: immunoglobulin variable-region peptides and the B-cell-receptor component CD79A, which are lower in autoimmune thyroid disease, and proteasome regulatory subunits together with the NF-κB subunit RELA, which are higher. A cross-validated classifier reproduced this separation (AUC 0.72, permutation p=0.05) with a stable eight-protein core, whereas within-class GHD severity was resolved by neither differential expression nor classification.

The biology is internally consistent. Autoimmune thyroid disease (Hashimoto’s and Graves’) is driven by B-cell and antibody responses against thyroid antigens, and the coordinated involvement of immunoglobulin/B-cell-receptor proteins alongside proteasome and NF-κB machinery — central to antigen processing and inflammatory transcription — is the expected molecular neighborhood for these conditions [6–10]. That the immunoglobulin variable-region signal was measured as lower in the autoimmune class in this particular panel, most markedly in Graves’ disease, is a quantitative observation about these specific peptides and should be interpreted in the context of a targeted assay rather than as a statement about total circulating immunoglobulin, whose titers are themselves an imperfect index of thyroid autoimmunity [11, 12]; it warrants targeted confirmation.

Equally important is what the panel could not do. The four groups were not globally separable, GHD severity was not predictable, and no protein reached FDR<0.05. Rather than obscure these negatives, we foreground them: they delineate the operating envelope of fixed-panel targeted proteomics in endocrinology. A panel weighted toward intracellular signaling, transcriptional, and proteasomal proteins captures an autoimmune-inflammatory axis well but is not positioned to track the metabolic gradations that distinguish partial from complete GH deficiency — gradations that remain difficult to resolve even with established clinical and biochemical testing [2, 4, 14]. The same principle — that a molecular classifier succeeds when the disease contrast aligns with the biology sampled by the assay — has been observed when plasma-proteomic classifiers separate clinically overlapping diseases with distinct immune signatures [16, 24].

Several constraints bound these conclusions. First, the autoimmune groups were small (Graves’ n=5, Hashimoto’s n=7), limiting statistical power; consistent with this, no protein survived FDR<0.05 and the class classifier reached only modest discrimination (AUC 0.72, permutation p=0.05 — at the significance boundary). Second, the panel is targeted and composition-biased toward signaling, transcription, and proteasome proteins, so it samples a limited region of the plasma proteome and cannot be read as proteome-wide. Third, growth-hormone-deficiency severity was not separable at the proteome level, consistent with the intrinsic difficulty of grading this axis even by established provocative testing [2, 14]. Fourth, no independent validation cohort was available, so all findings are hypothesis-generating. Fifth, clinical and biochemical covariates (age, sex, thyroid- and GH-axis hormone levels, medication, autoantibody titers) were not available for adjustment, and residual confounding cannot be excluded. Validation in a larger, clinically annotated cohort with orthogonal quantification of the immunoglobulin/B-cell-receptor and proteasome/NF-κB modules is the logical next step.

## Conclusion

Within a modest cohort and without external validation, a fixed 256-protein targeted panel reproducibly distinguishes autoimmune thyroid disease from growth-hormone deficiency along an immunoglobulin/B-cell-receptor and proteasome/NF-κB axis, while failing to resolve within-class severity. By reporting both the reproducible signal and its explicit limits, this study calibrates realistic expectations for targeted plasma-proteomic classification of endocrine disease and identifies concrete, testable markers for validation.

## Data and code availability

The mass spectrometry data have been deposited to Panorama Public (https://panoramaweb.org/), and are accessible at https://panoramaweb.org/co8XKz.url. All analysis scripts used in this study, including cosine similarity calculation, differential abundance analysis (limma), and figure generation, are publicly available on GitHub (https://github.com/kimlab-cnu/PedEndoMarker)

## Acknowledgement

The biospecimens and data used for this study were provided by the Biobank of Pusan National University Yangsan Hospital. All materials obtained (with informed consent) under institutional review board (IRB)-approved protocols. We acknowledge the use of Claude Opus 4.8 solely for linguistic refinement and grammatical corrections in manuscript preparation. All scientific content, data analysis, and intellectual contributions presented herein were developed independently by the authors without the use of generative AI tools.

## Author contributions

E.H. and J.J.: conceptualization, methodology, experimental analysis, data analysis, visualization; Y.C., J.P., H.L., S.M., and A.S.: data analysis; CKC and H.K.: writing-original draft, conceptualization, project administration, resources, supervision, writing-review & editing. All authors have read and approved the final manuscript.

